# The Network of Cancer Genes (NCG): a comprehensive catalogue of known and candidate cancer genes from cancer sequencing screens

**DOI:** 10.1101/389858

**Authors:** Dimitra Repana, Joel Nulsen, Lisa Dressler, Michele Bortolomeazzi, Santhilata Kuppili Venkata, Aikaterini Tourna, Anna Yakovleva, Tommaso Palmieri, Francesca D. Ciccarelli

**Author notes:** Equal contribution (Dimitra Repana, Joel Nulsen, Lisa Dressler, Michele Bortolomeazzi, Santhilata Kuppili Venkata).

## Abstract

The Network of Cancer Genes (NCG) is a manually curated repository of 2,372 genes whose somatic modifications have a known or predicted cancer driver role. These genes were collected from 275 publications, including two sources of known cancer genes and 273 cancer sequencing screens of 119 cancer types in 31 primary sites from 34,905 cancer donors. This represents a more than 1.5-fold increase in content as compared to the previous version. NCG also annotates properties of cancer genes, such as duplicability, evolutionary origin, RNA and protein expression, miRNA and protein interactions, protein function and essentiality. NCG is accessible at http://ncg.kcl.ac.uk/.

## BACKGROUND

One of the main goals of cancer genomics is to find the genes that, upon acquiring somatic alterations, play a role in driving cancer (cancer genes). To this end, in the last ten years, hundreds of cancer sequencing screens have generated mutational data from thousands of cancer samples. These include large sequencing efforts led by international consortia such as the International Cancer Genome Consortium (ICGC) [1] and The Cancer Genome Atlas (TCGA) [2]. Cancer genomes usually acquire thousands of somatic alterations and several methods have been developed to identify cancer genes from the pool of all altered genes [3, 4]. These methods have been applied to specific datasets from individual cancer types and to pooled datasets from several cancer types. This is the case for the Pan-Cancer Atlas project [5] and for the recent analysis of the whole set of TCGA samples [6], which accompanied the conclusion of the TCGA sequencing phase [7]. As more and more studies contribute to our knowledge of cancer genes, it becomes increasingly challenging for the research community to maintain an up-to-date overview of cancer genes and of the cancer types to which they contribute.

The Network of Cancer Genes (NCG) is a project launched in 2010 with the aim to gather a comprehensive and curated collection of cancer genes from cancer sequencing screens and to annotate their systems-level properties [8-11]. These define distinctive properties of cancer genes compared to other human genes [12] and include gene duplicability, evolutionary origin, RNA and protein expression, miRNA and protein interactions, protein function and essentiality. NCG is based on the manual curation of experts who review studies describing cancer sequencing screens, extract the genes that were annotated as cancer genes in the original publications and collect and analyse the supporting evidence.

Various other databases have been developed to analyse cancer data. Some of them focus on cancer alterations rather than cancer genes (COSMIC [13], DoCM [14], DriverDB [15], the Cancer Genome Interpreter [16], OncoKB [17], and cBIOPortal [18] among others). Other databases collect only cancer genes with a strong indication of involvement in cancer (the Cancer Gene Census, CGC [19]), annotate specifically oncogenes or tumour suppressor genes (ONGene [20], TSGene [21]) or cancer genes in specific cancer types (CoReCG [22]). NCG differs from all the above resources because it does not focus on mutations, on particular groups of genes or cancer types. It instead compiles a comprehensive repository of mutated genes that have been proven or predicted to be the drivers of cancer. NCG is widely used by the community. Recent examples of its use include studies identifying and validating cancer genes [23-25] and miRNA cancer biomarkers [26]. NCG has also been used to investigate the effect of long noncoding RNAs on cancer genes [27] and to find non-duplicated cancer-related transcription factors [28].

Here, we describe the sixth release of NCG, which contains 2,372 cancer genes extracted from 275 publications consisting of two sources of known cancer genes and 273 cancer sequencing screens. As well as mutational screens of individual cancer types, the collected publications now include four adult and two paediatric pan-cancer studies. In addition to an update of the systems-level properties of cancer genes already present in previous releases (gene duplicability, evolutionary origin, protein function, protein-protein and miRNA-target interactions, and mRNA expression in healthy tissues and cancer cell lines), NCG now also annotates the essentiality of cancer genes in human cell lines and their expression at the protein level in human tissues. Moreover, broader functional annotations of cancer genes in KEGG [29], Reactome [30] and BioCarta [31] are also provided.

The expert curation of a large number of cancer sequencing screens and the annotation of a wide variety of systems-level properties make NCG a comprehensive and unique resource for the study of genes that promote cancer.

## CONSTRUCTION AND CONTENT

The NCG database integrates information about genes with a known or predicted driver role in cancer. To facilitate the broad use of NCG, we have developed a user-friendly, interactive and open-access web portal for querying and visualising the annotation of cancer genes. User queries are processed interactively to produce results in a constant time. The front-end is connected to a database, developed using relational database management system principles [32] (Additional file 1: Figure S1). The web application for the NCG database was developed using MySQL v.5.6.38 and PHP v.7.0. Raw data for each of the systems-level properties were acquired from heterogeneous data sources and processed as described below. The entire content of NCG is freely available and can be downloaded from the database website.

### Gene duplicability and evolutionary origin

Protein sequences from RefSeq v.85 [33] were aligned to the human genome assembly hg38 with BLAT [34]. From the resulting genomic alignments, 19,549 unique gene loci were identified and genes sharing at least 60% of the original protein sequence were considered to be duplicated [35] (Additional file 2: Table S1). Orthologous genes for 18,486 human genes (including 2,348 cancer genes, Additional file 2: Table S1) in 2,032 species were collected from EggNOG v.4.5.1 [36] and used to trace the gene evolutionary origin as previously described [37]. Genes were considered to have a pre-metazoan origin if their orthologs could be found in prokaryotes, unicellular eukaryotes or opisthokonts [37].

### Gene and protein expression

RNA-Seq data from healthy human tissues for 18,984 human genes (including all 2,372 cancer genes, Additional file 2: Table S1) were derived from the non-redundant union of Protein Atlas v.18 [38] and GTEx v.7 [39]. Protein Atlas reported the average Transcripts Per Million (TPM) values in 37 tissues, and genes were considered to be expressed in a tissue if their expression value was ≥1 TPM. GTEx reported the distribution of TPM values for individual genes in 11,688 samples across 30 tissue types. In this case, genes were considered to be expressed if they had a median expression value ≥1 TPM.

Gene expression data for all 2,372 cancer genes in 1,561 cancer cell lines were taken from the Cancer Cell Line Encyclopedia (CCLE, 02/2018) [40], the COSMIC Cancer Cell Line Project (CLP, v.84) [19] and a Genentech study (GNE, 06/2014) [41] (Additional file 2: Table S1). Gene expression levels were derived directly from the original sources, namely Reads Per Kilobase Million (RPKM) values for CCLE and GNE, and microarray z-scores for CLP. Genes were categorised as expressed if their expression value was ≥1 RPKM in CCLE or GNE, and were annotated as over, under or normally expressed in CLP, as determined by COSMIC.

The current release of NCG also includes protein expression from immunohistochemistry assays of healthy human tissues as derived from Protein Atlas v.18. Data were available for 13,001 human proteins including 1,799 cancer proteins (Additional file 2: Table S1). Proteins were categorised as not detected or as having low, medium or high expression in 44 tissues on the basis of staining intensity and fraction of stained cells [38]. In Protein Atlas, expression levels were reported in multiple cell types for each tissue. NCG retained the highest reported value as the expression level for that tissue.

### Gene essentiality

Gene essentiality was derived from two databases, PICKLES (09/2017) [42] and OGEE v.2 [43], both of which collected data from CRISPR Cas9 knockout and shRNA knockdown screens of human cell lines. In PICKLES, data from primary publications have been re-analysed and genes were considered essential in a cell line if their associated Bayes factor was >3 [44]. We therefore used this threshold to define essential genes. In OGEE, genes were labelled as “essential” or “not essential” according to their annotation in the original publications. Consistently, we retained the same annotation. From the non-redundant union of the two databases, essentiality information was available for a total of 18,833 genes (including all 2,372 cancer genes) in 178 cell lines (Additional file 2: Table S1).

### Protein-protein and miRNA-target interactions

Human protein-protein interactions were derived from four databases (BioGRID v.3.4.157 [45]; MIntAct v.4.2.10 [46]; DIP (02/2018) [47] and HPRD v.9 [48]). Only interactions between human proteins supported by at least one original publication were considered [8]. The union of all interactions from the four sources was used to derive a human protein-protein interaction network of 16,322 proteins (including 2,203 cancer proteins, Additional file 2: Table S1) and 289,368 binary interactions. To control for a possible higher number of studies on cancer proteins resulting in an artificially higher number of interactions, a network of 15,272 proteins and 224,258 interactions was derived from high-throughput screens reporting more than 100 interactions [11].

Data on human protein complexes for 8,080 human proteins (including 1,414 cancer proteins, Additional file 2: Table S1) were derived from the non-redundant union of three primary sources, namely CORUM (07/2017) [49], HPRD v.9 [48] and Reactome v.63 [30]. Only human complexes supported by at least one original publication were considered [11].

Experimentally validated interactions between human genes and miRNAs were downloaded from miRTarBase v.7.0 [50] and miRecords v.4.0 [51], resulting in a total of 14,649 genes (including 2,034 cancer genes) and 1,762 unique miRNAs (Additional file 2: Table S1). To control for the higher number of single-gene studies focussing on cancer genes, a dataset of high-throughput screens testing ≥250 different miRNAs was also derived (Additional file 2: Table S1).

### Functional annotation

Data on functional categories (pathways) were collected from Reactome v.63 [30], KEGG v.85.1 [29] and BioCarta (02/2018) [31]. Data for BioCarta were extracted from the Cancer Genome Anatomy Project [52]. All levels of Reactome were included, and level 1 and 2 pathways from KEGG were added separately. Overall, functional annotations were available for 11,344 human proteins, including 1,750 cancer proteins assigned to 2,318 pathways in total.

## UTILITY AND DISCUSSION

### Catalogue of known and candidate cancer genes

To include new cancer genes in NCG, we applied a modified version of our well-established curation pipeline [11] (Figure 1A). We considered two main groups of cancer genes: known cancer genes whose involvement in cancer has additional experimental support, and candidate cancer genes whose somatic alterations have a predicted cancer driver role but lack further experimental support.

**Figure 1.**
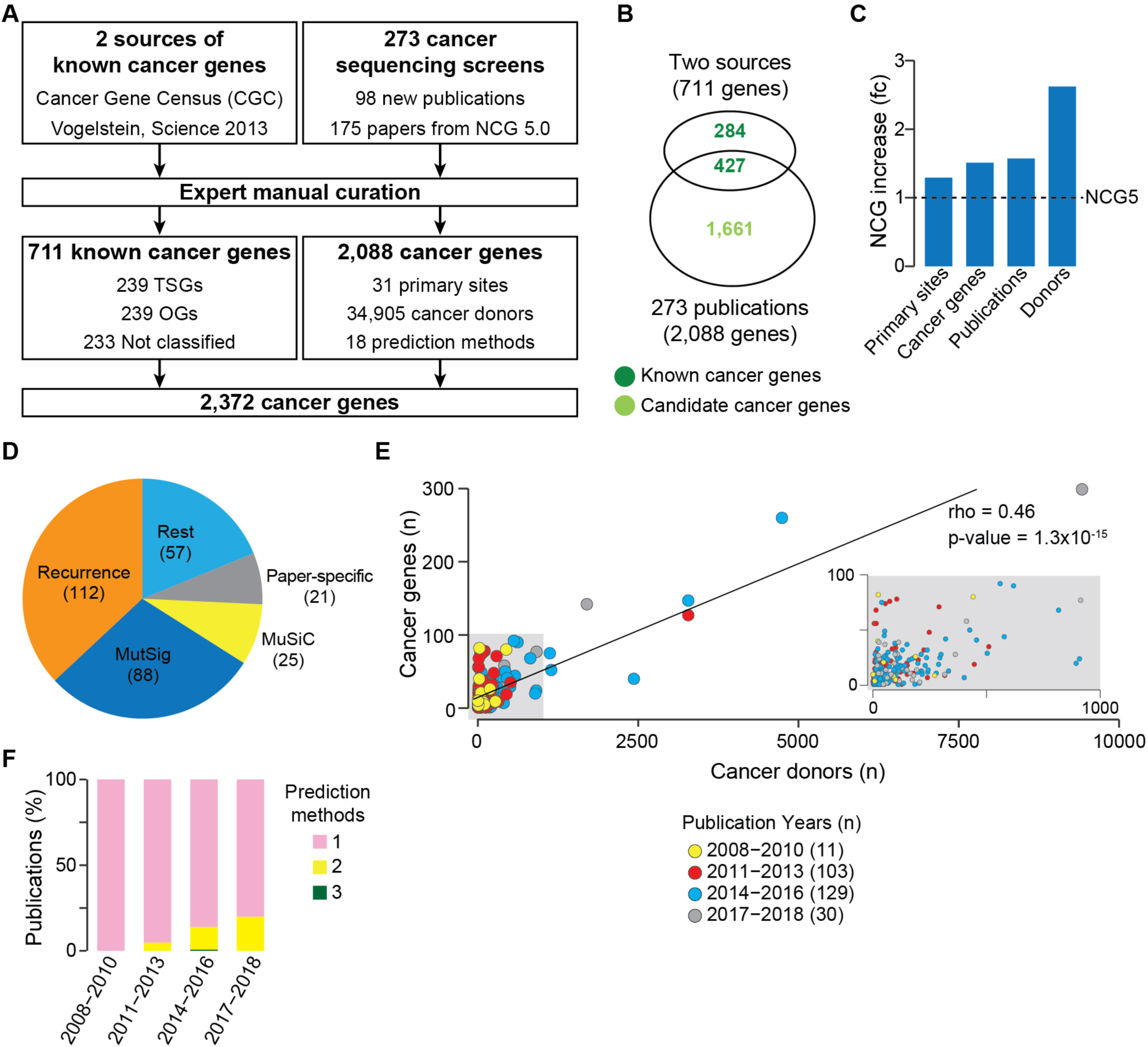
Manual curation of cancer genes in NCG. **a.** Pipeline used for adding cancer genes to NCG. Two sources of known cancer genes [19, 53] were integrated leading to 711 known cancer genes. In parallel, 273 publications describing cancer sequencing screens were reviewed to extract 2,088 cancer genes. The non-redundant union of these two sets led to 2,372 cancer genes currently annotated in NCG. **b.** Intersection between known and candidate cancer genes in NCG. **c.** Comparison of NCG content with the previous version [11]. **d.** Pie chart of the methods used to identify cancer genes in the 273 publications. The total is greater than 273 because some studies used more than one method (Additional file 2: Table S2). **e.** Cancer genes as a function of the number of cancer donors per study. The grey inset shows a magnification of the left bottom corner of the plot. **f.** Number of methods used to identify cancer genes over time. PanSoftware used in one of the pan-cancer studies [6] was considered as a single method but is in fact a combination of 26 prediction tools.

As sources of known cancer genes, we used 708 genes from CGC v.84 [19] and 125 genes from a manually curated list [53]. Of the resulting 711 genes, we further annotated 239 as tumour suppressor genes (TSGs) and 239 as oncogenes (OGs). The remaining 233 genes could not be unambiguously classified because either they had conflicting annotations in the two original sources (CGC and [53]) or they were defined as both OGs and TSGs. Despite these two sources of known cancer genes have been extensively curated, 49 known cancer genes are in two lists of possible false positives [6, 54].

Next, we reviewed the literature to search for studies that (1) described sequencing screens of human cancers and (2) provided a list of genes considered to be the cancer drivers. This led to 273 original papers published between 2008 and March 2018, 98 of which were published since the previous release of NCG [11] and 42 of which came from ICGC or TGCA (Additional file 2: Table S2). Overall, these publications describe the sequencing screens of 119 cancer types from 31 primary anatomical sites as well as six pan-cancer studies (Additional file 2: Table S2). In total, this amounts to samples from 34,905 cancer donors. Each publication was reviewed independently by at least two experts and all studies whose annotation differed between the experts were further discussed. Additionally, 31 randomly selected studies (11% of the total) were re-annotated blindly by a third expert to assess consistency. The manual revision of the 273 studies led to 2,088 cancer genes, of which 427 were known cancer genes and the remaining 1,661 were candidate cancer genes (Figure 1B). Compared to the previous release, this version of NCG constitutes a significant increase in the number of cancer primary sites (1.3-fold), cancer genes (1.5-fold), publications (1.6-fold), and analysed donors (2.6-fold, Figure 1C).

Based on literature evidence [6, 54], gene length and function [10], 201 candidates were labelled as possible false positive predictions. We further investigated the reasons why 284 known cancer genes were not identified as drivers in any of the 273 cancer sequencing screens. We found that these genes predispose to cancer rather than acquiring somatic alterations, are the chimeric product of gene fusions, are part of CGC Tier 2 (*i.e.* genes with lower support for their involvement in cancer) or were identified with different methods. Eleven of these 284 genes are possible false positives [6, 54].

The annotation of a large number of studies allowed us to gain insights into how cancer genes have been identified in the last ten years. Of the overall 18 prediction methods (Additional file 2: Table S2), the recurrence of a gene alteration within the cohort is the most widely used across screens (Figure 1D). In this case, no further threshold of statistical significance or correction for the genome, gene and cancer background mutation rate was applied, thus leading to possible false positive predictions. Other frequently used prediction methods are MutSig [55], MuSiC [56] and *ad hoc* pipelines developed in the same publication (referred as ‘paper-specific’). Although they apply statistical methods to correct for the background mutation rate and reduce false positives, all of these approaches estimate the tendency of a gene to mutate more than expected within a cohort and therefore they all depend on sample size. Indeed, we observed an overall positive correlation between the number of cancer donors and the number of cancer genes (Figure 1E). This confirms that the sensitivity of the approaches currently used to predict cancer genes is higher for large cohorts of samples. Finally, although the vast majority of analysed studies tend to apply only one prediction method, more recent publications have started to use a combination of two or three methods (Figure 1F).

### Heterogeneity and specificity of cancer genes

The number of cancer genes and the relative proportion of known and candidate cancer genes vary greatly across cancer primary sites (Figure 2A). More than 75% of cancer genes in cancers of the prostate, soft tissues, bone, ovary, cervix, thymus and retina are known drivers. On the contrary, more than 75% of driver genes in cancers of the penis, testis and vascular system are candidate cancer genes (Figure 2A). This seems to be due to several factors including the sample size, the number of different methods that have been applied to identify cancer genes and the biology of each cancer type. For example, penis, vascular system and testis cancers show a high proportion of candidate cancer genes. The corresponding cohorts have a small sample size and have been analysed by one or two prediction methods. However, other cancer types showing equally high proportions of candidates (pancreas, skin, blood) have large sample sizes and were analysed by several methods (Figure 2B). Moreover, although the number of cancer genes is overall positively correlated with the number of sequenced samples (Figures 1C, 2C), there are marked differences across primary sites. For example, ovary, bone, prostate, thyroid and kidney cancers have substantially fewer cancer genes compared to cancers with similar numbers of cancer donors such as uterine, stomach, skin and hepatobiliary cancers (Figure 2C). This is likely due to variable levels of genomic instability and heterogeneity across cancer types of the same primary site. For example, in seven of the nine mutational screens of skin melanoma, a cancer type with high genomic instability [57], more than 50% of cancer genes are study-specific (Figure 3A). Similarly, the 24 types of blood cancer are variable in terms of number of cancer genes, with diffuse large B-cell lymphoma having many more cancer genes than other blood cancers with higher numbers of cancer donors (Figure 3B). In both cases, the use of the same method (i.e. MutSig in Figure 3A and MuSiC in Figure 3B) identified different cancer genes in different patient cohorts, highlighting the biological heterogeneity even across donors of the same cancer type.

**Figure 2.**
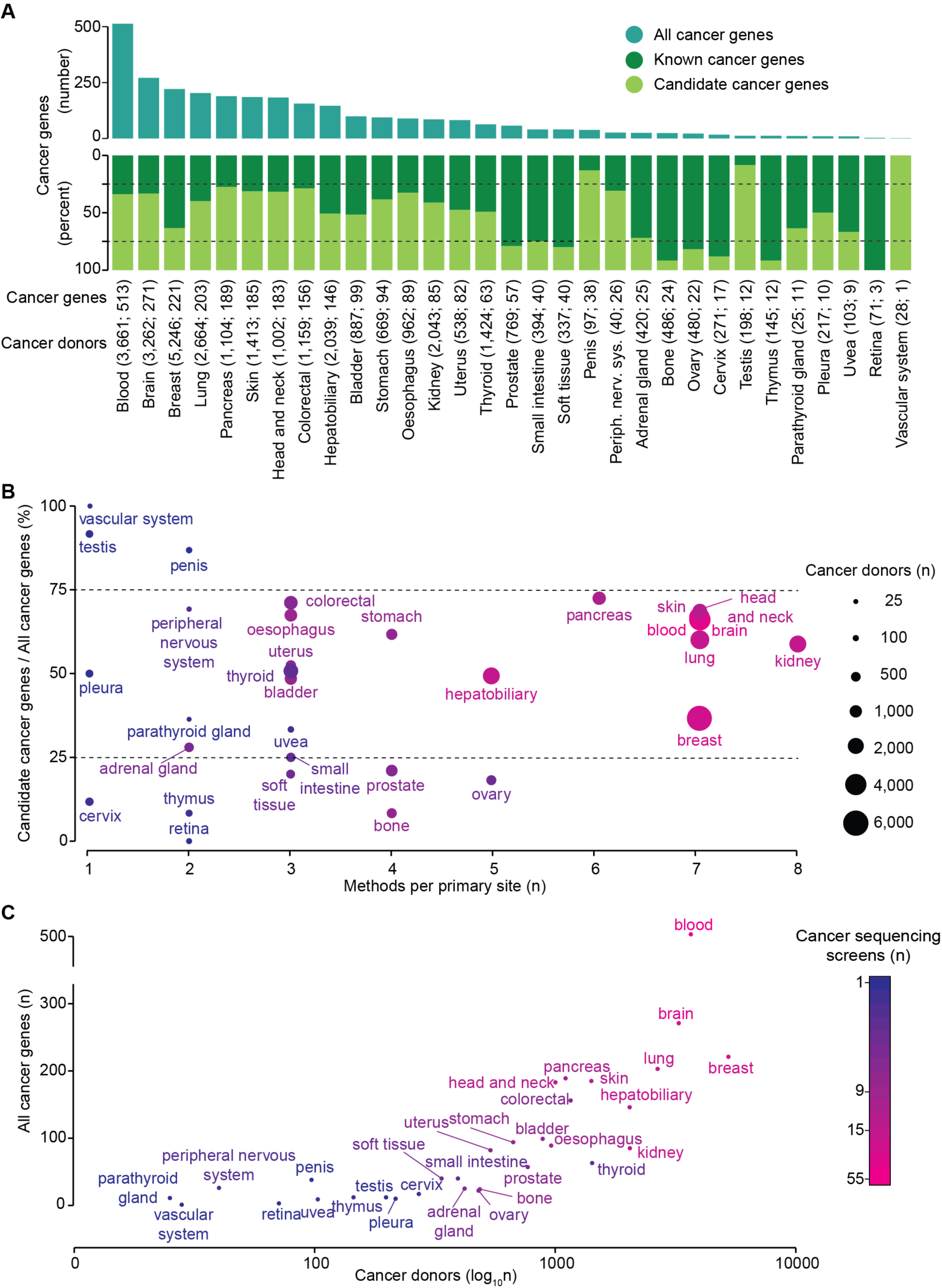
Distribution of cancer genes across primary sites and cancer donors. **a.** Number of total cancer genes and proportion of known and candidate cancer genes across the 31 tumour primary sites analysed in the 267 cancer-specific studies. The number of cancer donors followed by the number of cancer genes is given in brackets for each primary site. **b.** Proportion of candidate cancer genes over all cancer genes across the 31 tumour primary sites. The dot size is proportional to the donor cohort size. **c.** Total number of cancer genes and cancer donors across the 31 tumour primary sites. The colour scale in (**b**) and (**c**) indicates the number of screens for each primary site.

**Figure 3.**
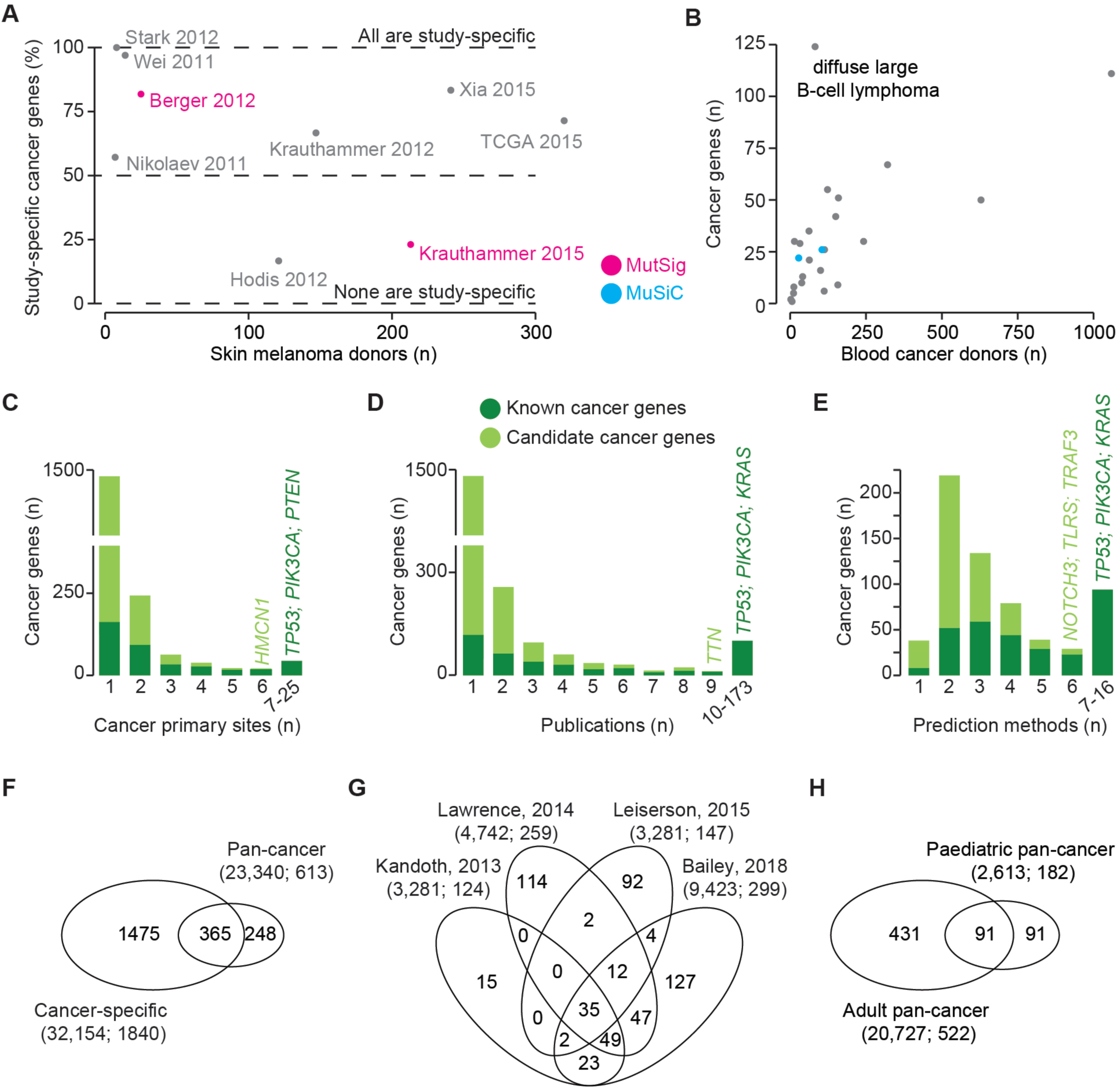
Recurrence of cancer across primary sites and publications. **a.** Proportion of study-specific cancer genes reported by each of the seven skin melanoma screens. **b.** Total number of cancer genes and donors across 24 cancer types of the blood. The full list of blood cancer types is reported in Additional file 2: Table S2. **c.** Number of primary sites in which each known or candidate cancer gene was reported to be a driver. **d.** Number of publications in which each known or candidate cancer gene was reported to be a driver. **e.** Number of methods used to predict cancer genes for drivers found in more than one publication. **f.** Intersection of cancer genes in the cancer-specific and pan-cancer studies. **g.** Venn diagram of cancer genes across the four pan-cancer studies of adult donors. **h.** Intersection of cancer genes in pan-cancer screens of adult and paediatric donors. In **f**, **g**, and **h** the number of donors followed by the total number of cancer genes are given in brackets.

Cancer genes, and in particular candidates, are highly cancer specific (Figure 3C). Hemicentin 1 (*HMCN1*) is the only candidate cancer gene to be significantly mutated in six primary sites (blood, brain, oesophagus, large intestine, liver, and pancreas). A few known cancer genes are recurrently mutated across several primary sites, including *TP53* (25), *PIK3CA* (21) and *PTEN* (20, Figure 3C). These are, however, exceptions and the large majority of known and candidate cancer genes (64% of the total) are found only in one primary site, indicating high heterogeneity of cancer driver events. Similar specificity is also observed in terms of supporting publications. The majority of cancer genes are publication-specific, again with few exceptions including *TP53* (173), *PIK3CA* (87) and *KRAS* (86, Figure 3D). Of note, the best-supported candidate gene is Titin (*TTN*, predicted in nine publications), which is a well-known false positive of recurrence-based approaches [55]. Interestingly, the scenario is different when analysing the number of prediction methods that support cancer genes reported in at least two screens (Figure 3E). In this case, few candidate and known cancer genes are identified by only one method, while the majority of them are supported by at least two (candidates) and three (known cancer genes) approaches. However, only 6 candidate cancer genes are supported by six methods and *TP53* is the only cancer genes to be identified by 16 of the 18 methods (Figure 3E).

Finally, the heterogeneity of the cancer driver landscape is reflected in the pan-cancer studies. Approximately 40% of the cancer genes from pan-cancer analyses were not previously predicted as drivers (Figure 3F), despite the large majority of cancer samples having been already analysed in the corresponding cancer-specific study. This is yet a further confirmation that current methods depend on the sample size and that a larger cohort leads to novel predictions. Only 35 cancer genes were shared across four pan-cancer re-analyses of adult tumours (Figure 3G), suggesting that the prediction of cancer genes is highly method-and cohort-dependent. This is further confirmed by the poor overlap between cancer genes from adult and paediatric pan-cancer studies (Figure 3H). In this case, however, it is also likely that different biological mechanisms are responsible for adult and childhood cancers.

Overall, our analysis of the cancer driver landscape suggests that the high heterogeneity of cancer genes observed across cancer types is due to a combination of sample size effect, prediction methods and true biological differences across cancers.

### Systems-level properties of cancer genes

In addition to collecting cancer genes from the literature, NCG also annotates the systems-level properties that distinguish cancer genes from other genes that are not implicated in cancer (Additional file 2: Table S1). We therefore compared each of these properties between cancer genes and the rest of human genes. We considered seven distinct groups of cancer genes. The first three were 711 known cancer genes, 1,661 candidate cancer genes and 2,372 total cancer genes. After removing 201 possible false positive predictions [6, 54] from the list of candidate cancer genes, we also identified two sets of candidate cancer genes with a stronger support. One was composed of 104 candidate cancer genes found in at least two independent screens of the same primary site. The other was formed of 711 candidate cancer genes identified in large cohorts composed of at least 140 donors (top 25% of the sample size distribution across screens). Finally, we compared the properties between 239 TSGs and 239 OGs.

As previously reported [35], we confirmed that a significantly lower fraction of cancer genes has duplicated copies in the human genome due to a high proportion of single-copied TSGs (Figure 4A). The same trend was observed in both known and candidate cancer genes, and is significant for the combination of the two gene sets. Interestingly, candidate cancer genes found in ≥ 2 screens show a high proportion of duplicated cancer genes (albeit not significant probably due to the small size of the group, Figure 2B). This could suggest that several genes in this group may exert an oncogenic role.

**Figure 4.**
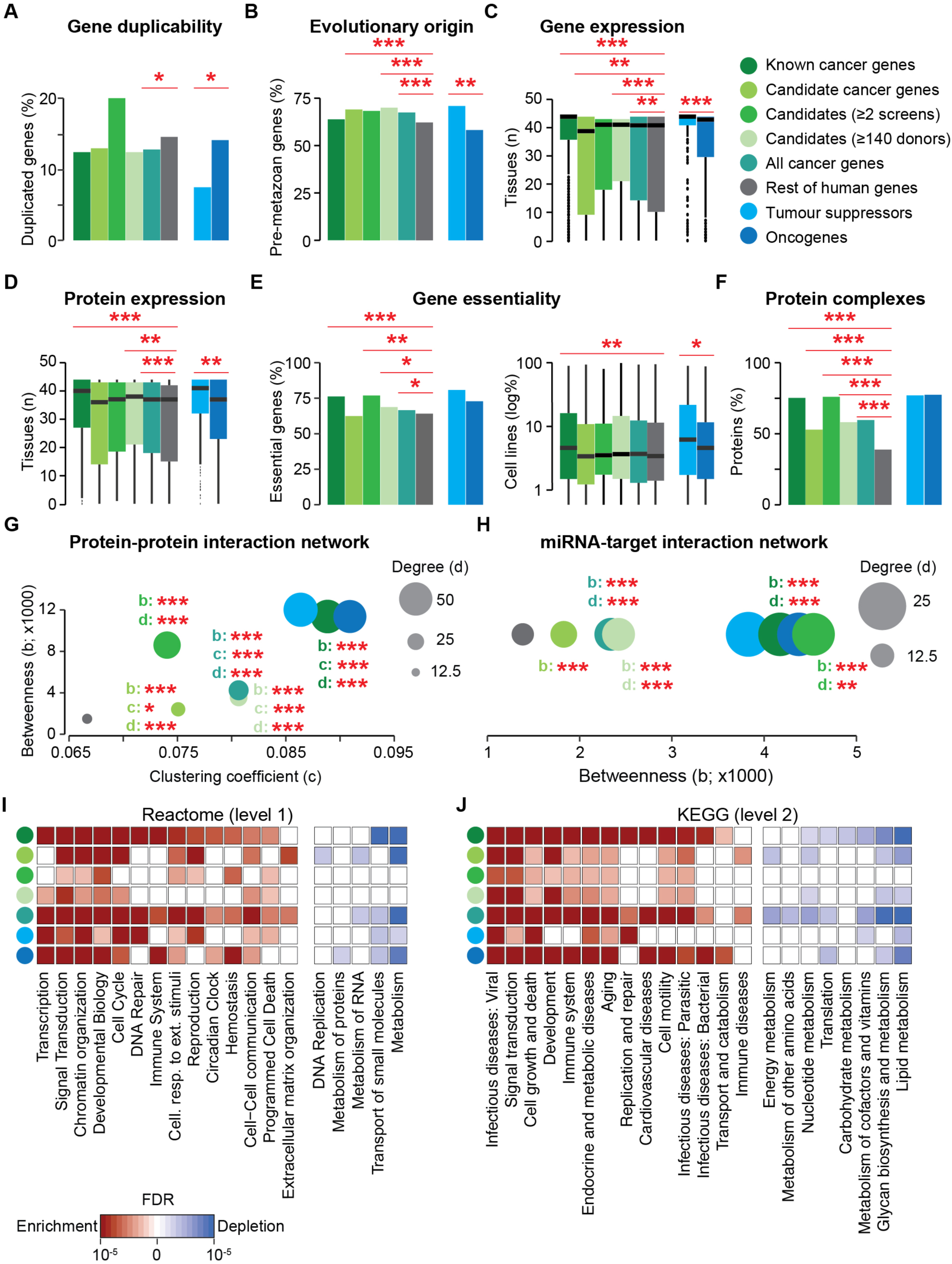
Systems-level properties of cancer genes. **a.** Percentage of genes with ≥1 gene duplicate covering ≥60% of the protein sequence. **b.** Proportion of genes originating in pre-metazoan species. **c,d.** Number of human tissues in which genes (**c**) and proteins (**d**) are expressed. In panel **c**, tissue types were matched between GTEx and Protein Atlas wherever possible, giving 43 unique tissues. In tissues represented in both datasets, genes were defined as expressed if they had ≥1 TPM in both datasets. Only genes present in both sources were compared (Additional file 2: Table S1). **e.** Percentage of genes essential in ≥1 cell line and distribution of cell lines in which each gene is essential. Only genes with concordant annotation between OGEE and PICKLES were compared (Additional file 2: Table S1). **f.** Percentage of proteins involved in ≥1 protein complex. **g.** Median values of betweenness (centrality), clustering coefficient (clustering) and degree (connectivity) of human proteins in the protein-protein interaction network. **h.** Median values of betweenness and degree of the target genes in the miRNA-target interaction network. The clustering coefficient is zero for all nodes, because interactions occur between miRNAs and target genes. Known, candidate and all cancer genes were compared to the rest of human genes, while TSGs were compared to OGs. Significance was calculated using a two-sided Fisher test (**a,b,e,f**) or Wilcoxon test (**c,d,g,h**). * = p<0.05, ** = p<0.01, *** = p<0.001. Enrichment and depletion of cancer genes in representative functional categories taken from level 1 of Reactome (**i**) and level 2 of KEGG (**j**). Significance was calculated comparing each group of cancer genes to the rest of human genes using a two-sided Fisher test. False discovery rates were calculated in each gene set separately. Only pathways showing enrichment or depletion are shown. The full list of pathways is provided in Additional file 2: Table S3.

Cancer genes, and in particular candidate cancer genes, originated earlier in evolution (Figure 4B) [37, 58, 59]. Known cancer genes alone do not differ from the rest due to the fact that OGs are significantly younger than TSGs (Figure 4B).

Known cancer genes tend to be ubiquitously expressed at the mRNA (Figure 4C) and protein (Figure 4D) levels and TSGs are more widely expressed than OGs. This trend is less clear when analysing candidate cancer genes separately. Candidates with stronger support tend to resemble known cancer genes, however the overall set of candidate cancer genes has a narrower tissue expression pattern at the gene and protein level (Figure 4C, D)

A similar scenario is observed when analysing gene essentiality. A higher fraction of cancer genes, and in particular of known cancer genes, is essential in at least one human cell line (Figure 4E). Moreover, known cancer genes tend to be essential in a higher fraction of cell lines. Both measures of gene essentiality are higher in TSGs as compared to OGs (Figure 4E). Candidate cancer genes with stronger support are again similar to known cancer genes but, when considered together, all candidate cancer genes are not significantly enriched in essential genes (Figure 4E).

Proteins encoded by cancer genes are more often involved in protein complexes (Figure 4F). They are also more connected (higher degree), central (higher betweenness) and clustered (higher clustering coefficient) in the protein-protein interaction network (Figure 4G). We verified that this trend holds true also when using only data from high-throughput screens (Additional file 2: Table S2), thus excluding the possibility that the distinctive network properties of cancer proteins are due to their better annotation. These trends remain significant for all sets of cancer genes.

Cancer genes are regulated by a higher number of miRNAs (higher degree) and occupy more central positions (higher betweenness) in the miRNA-target interaction network (Figure 4H). As above, these results remain valid also when only considering the miRNA-target network from high throughput screens (Additional file 2: Table S2) and for any group of cancer genes considered.

Cancer genes are consistently enriched in functional categories such as signal transduction, chromatin reorganisation and cell cycle and depleted in others, such as metabolism and transport (Figure 4I, Additional file 2: Table S3). Candidate cancer genes generally exhibit weaker enrichment than the other groups, most notably in DNA repair. Interestingly, however, extracellular matrix reorganisation displays a specific enrichment for candidate cancer genes. Some functional categories are selectively enriched for OGs (*i.e.* development, immune and endocrine systems, Figure 4J) or TSGs (*i.e.* DNA repair and programmed cell death). While annotations from Reactome and KEGG generally give concordant results, they differ significantly for gene transcription. In this case, Reactome shows a strong enrichment for cancer genes, while it is not significant in KEGG (Figure 4I,J).

Overall our analyses confirm that cancer genes are a distinctive group of human genes. Despite their heterogeneity across cancer types and donors, they share common properties. Candidate cancer genes only share some of the properties of known cancer genes, such as an early evolutionary origin (Figure 4B) and higher centrality and connectivity in the protein-protein and miRNA-target interaction networks (Figure 4G, H). They do not differ from the rest of genes for all other properties. However, the two sets of candidate cancer genes with a stronger support overall maintain the vast majority of the distinctive properties of known cancer genes. This suggests that the current set of candidate cancer genes likely contains false positives and genes with weak support that do not resemble the properties of known cancer genes. This is further indicated when directly comparing the properties of known and candidate cancer genes (Additional file 2: Table S4). In this case, known cancer genes are significantly different for most properties when compared to the whole set of candidate cancer genes. However, these differences are reduced when he two sets of candidates with stronger support are used. Finally, TSGs and OGs constitute two distinct classes of cancer genes even based on their systems-level properties (Figure 4).

### Future directions

In the coming years, NCG will continue to collect new cancer genes and annotate their properties, including novel properties such as genetic interactions or epigenetic features for which large datasets are becoming available. So far, the cancer genomics community has focussed mostly on the identification of protein-coding genes with putative cancer driver activity. With the increasing availability of whole genome sequencing data and a rising interest in non-coding alterations [27, 60], NCG will expand to also collect non-coding cancer drivers. Another direction for future development will be the analysis of clinical data, including therapeutic treatments, to link them to the altered drivers. This will contribute to the expansion of our knowledge of cancer driver genes in the context of their clinical relevance.

## CONCLUSIONS

The present release of NCG describes a substantial advance in annotations of known and candidate cancer driver genes as well as an update and expansion of their systems-level properties. The extensive body of literature evidence collected in NCG enabled a systematic analysis of the methods used to identify cancer genes, highlighting their dependence on the number of cancer donors. We also confirmed the high heterogeneity of cancer genes within and across cancer types. The broad set of systems-level properties collected in NCG shows that cancer genes form a distinct group, different from the rest of human genes. For some of these properties, the differences observed for known cancer genes hold true also for candidate cancer genes, and TSGs show more pronounced cancer gene properties than OGs. Interestingly, these properties are shared by all cancer genes, independently of the cancer type or gene function. Therefore, focussing on genes with similar characteristics could be used for the identification and prioritisation of new cancer driver genes [61]. In conclusion, the large-scale annotation of the systems-level properties of cancer genes in NCG is a valuable source of information not only for the study of individual genes, but also for the characterisation of cancer genes as a group.

## LIST OF ABBREVIATIONS

CCLE: Cancer Cell Line Encyclopedia
CLP: Cell Line Project (COSMIC)
GNE: Genentech
ICGC: International Cancer Genome Consortium
miRNA: microRNA
NCG: Network of Cancer Genes
OG: Oncogene
RPKM: Reads Per Kilobase Million
TCGA: The Cancer Genome Atlas
TPM: Transcripts Per Million
TSG: Tumour Suppressor Gene.

## DECLARATIONS

### 1 ETHICS APPROVAL AND CONSENT TO PARTICIPATE

Not applicable

### 2 CONSENT FOR PUBLICATION

Not applicable

### 3 AVAILABILITY OF DATA AND MATERIALS

The whole content of NCG can be downloaded from the website. Original data were obtained from the following online sources:

BioCarta: https://cgap.nci.nih.gov/Pathways/BioCarta_Pathways;

BioGRID: https://thebiogrid.org/;

Cancer Cell Line Encyclopedia: https://portals.broadinstitute.org/ccle;

Cell Line Project: https://cancer.sanger.ac.uk/cell_lines;

CORUM: http://mips.helmholtz-muenchen.de/corum/;

DIP: http://dip.doe-mbi.ucla.edu/dip/Main.cgi;

EggNOG: http://eggnogdb.embl.de/#/app/home;

Genentech: https://www.ebi.ac.uk/arrayexpress/experiments/E-MTAB-2706/;

GTEx: https://www.gtexportal.org/home/;

HPRD:http://www.hprd.org/;ICGC,https://icgc.org/;

KEGG: http://www.genome.jp/kegg/pathway.html;

MIntAct: https://www.ebi.ac.uk/intact/;

miRecrods:http://c1.accurascience.com/miRecords/;

miRTarBase: http://mirtarbase.mbc.nctu.edu.tw/php/index.php; OGEE:http://ogee.medgenius.info/browse/;

PICKLES: https://hartlab.shinyapps.io/pickles/;

Protein Atlas: https://www.proteinatlas.org/;

Reactome: https://reactome.org/;

RefSeq: https://www.ncbi.nlm.nih.gov/refseq/;

TCGA, https://cancergenome.nih.gov/

### 4 COMPETING INTERESTS

The authors declare that they have no competing interests.

### 5 FUNDING

This work was supported by Cancer Research UK [C43634/A25487] and by the Cancer Research UK King’s Health Partners Centre at King’s College London. Computational analyses were done using the High-Performance Computer at the National Institute for Health Research (NIHR) Biomedical Research Centre based at Guy’s and St Thomas’ NHS Foundation Trust and King’s College London. JN is supported by the EPSRC Centre for Doctoral Training in Cross-Disciplinary Approaches to Non-Equilibrium Systems (CANES, EP/L015854/1). DR is supported by the Health Education England (HEE) Genomics Education Programme (GEP).

### 6 AUTHORS’ CONTRIBUTIONS

FDC conceived and supervised the study. SKV, DR, JN, LD, MB, AT, and FDC analysed the data. SKV analysed gene duplicability. MB processed evolutionary origins and miRNA-target interactions. LD processed protein-protein interactions, protein complexes and gene essentiality. JN processed RNA and protein expression and protein function. DR, AT, AY and TP curated the literature. SKV and JN updated the database and website. JN, LD, MB, AT, and FDC wrote the manuscript, with contributions from SKV and DR. All authors reviewed and approved the final version of the manuscript.

## 7 ACKNOWLEDGMENTS

We thank all members of the Ciccarelli lab for useful feedback on the website and Elizabeth Foxall for proofreading of the manuscript.

## LIST OF SUPPLEMENTARY DATA

### Additional file 1 (JPEG)

**Figure S1. Schema of the NCG database**

Entity-relationship diagram indicating one-to-many and many-to-many relationships between genes and other entities in the NCG database. The external source files used to generate the Genes entity are shown in grey.

**Additional file 2 (XLSX)**

**Table S1. Systems-level properties of cancer genes.** For each systems-level property reported are the numbers of genes for which data were available, the percentage of annotated genes out of all the genes in each group and the p-values of the comparisons between groups (cancer genes versus the rest of human genes and tumour suppressor genes versus oncogenes). Duplicated genes (genes sharing at least 60% of the original protein sequence length) were compared between groups using a two-tailed Fisher test. The fractions of pre-metazoan genes in each group were compared with a chi-square test. For gene expression data, only genes with concordant annotations between original sources were compared between groups using two-tailed Wilcoxon tests. Similarly, only genes with concordant essentiality annotations in each cell line between the original sources were considered. The percentages of essential genes were compared using a two-tailed Fisher test. The distributions of cell lines in which genes are essential were compared using a two-tailed Wilcoxon test. The fraction of genes participating in at least one complex was compared using a two-tailed Fisher test. Distributions of degree, betweenness and clustering coefficient of proteins in the protein-protein interaction network were compared using a two-tailed Wilcoxon test. Distributions of degree and betweenness of genes in the miRNA-target interaction network were compared using a two-tailed Wilcoxon test. In this case, the clustering coefficient could not be measured because the interactions occur only between miRNAs and target genes. Versions of the database, references to website and details on the methods are given in the text and in the legend to Figure 4. TSG, tumour suppressor gene; OG, oncogene.

**Table S2. List of 273 publications describing cancer sequencing screens in NCG.** For each publication, provided are the number of cancer genes and cancer donors, the cancer primary site, the cancer type as reported in the publication, the methods used to identify cancer types, a description of the method and whether the study is part of the TCGA or ICGC consortia. Recurrence indicates that a gene was considered as a cancer driver if it was mutated recurrently within a cohort, without applying any further statistical threshold. Literature indicates that a gene was considered as a cancer driver based on literature evidence (it was a known cancer gene or it had been previously associated with cancer). Paper-specific methods indicate specific criteria defined by the authors in the corresponding publication.

**Table S3. Enrichment and depletion of cancer genes in protein functional categories.** Cancer genes were analysed for enrichment/depletion in functional categories (pathways) from Level 1 of Reactome v.63 or Level 2 of KEGG v.85.1. Seven groups of cancer genes (known cancer genes, candidate cancer genes, candidates reported by ≥2 screens in the same primary site, candidates in screens with ≥140 donors, all cancer genes, tumour suppressor genes (TSG) and oncogenes (OG) were compared to non-cancer genes with a two-sided Fisher test. The resulting p-values were corrected separately for different pathway sources and different comparisons to obtain the false discovery rate (FDR). Where a significant signal was found (FDR<0.05), the odds ratio being greater than or less than 1 was used to determine whether the signal was one of enrichment or depletion, respectively.

**Table S4. Comparison of systems-level properties between known and candidate cancer genes.** Each systems-level property was compared between known cancer genes and three groups of candidate cancer genes: all candidate cancer genes, candidates found in at least two screens of the same cancer type and candidates predicted in large cohorts (at least 140 donors, corresponding to the upper quartile of the distribution of cohort size across all screens). Properties were defined and compared as described in Supplementary Table S1.

